# Finite-sites multiple mutations interference gives rise to wavelet-like oscillations of multilocus linkage disequilibrium

**DOI:** 10.1101/090571

**Authors:** Victor Garcia, Emily C. Glassberg, Arbel Harpak, Marcus W. Feldman

**Affiliations:** Department of Biology, Stanford University, 371 Serra Mall, Stanford CA-94305, USA

## Abstract

Within-host adaptation of pathogens such as human immunodeficiency virus (HIV) often occurs at more than two loci. Multiple beneficial mutations may arise simultaneously on different genetic backgrounds and interfere, affecting each other's fixation trajectories. Here, we explore how these adaptive dynamics are mirrored in multilocus linkage disequilibrium (MLD), a measure of multi-way associations between alleles. In the parameter regime corresponding to HIV, we show that deterministic early infection models induce MLD to oscillate over time in a wavelet-like fashion. We find that the frequency of these oscillations is proportional to the rate of adaptation. This signature is robust to drift, but can be eroded by high variation in fitness effects of beneficial mutations. Our findings suggest that MLD oscillations could be used as a signature of interference among multiple equally advantageous mutations and may aid the interpretation of MLD in data.

## INTRODUCTION

Many microorganisms, viruses, and cancer cells replicate asexually with large population sizes and under strong selection [1–7]. This gives rise to pronounced genetic *interference* [2, 6, 8, 9], where beneficial mutations can emerge on different haplotypes and compete, leading to mutual growth impairment [10–16]. Since interference determines how asexual organisms adapt, it is of particular relevance to understanding infectious disease agents.

Most recent theoretical treatments of interference rest upon the assumption that there exists an infinite supply of new, beneficial mutations arising from an infinite number of loci [2, 16–19]. Indeed, recent studies in yeast and other microorganisms suggest that beneficial mutation supply is not what limits the rate of adaptation [1–6]. However, the infinite-sites assumption might be inappropriate for understanding many asexual populations evolving over short time scales under strong selective pressure. In fact, most short-term adaptation of pathogens to new host environments occurs at a limited number of either known [7], or detectable loci [20]. Key examples are drug resistance mutations or escape mutations in viruses [21], such as HIV [8, 22]. Furthermore, the selective pressures exerted on pathogens by a finite number of immune responses during early infection typically far exceed what other stresses might shape their adaptation [20, 22–25].

Past research on finite-sites interference typically involved only a few, predominantly two loci [12, 26–31]. Traditionally, pair-wise linkage disequilibrium (LD) has been a successful summary statistic for such interference [31, 32]. For example, if two mutations are physically linked by the same background, this will result in pronounced positive LD. If they disproportionately often appear on different backgrounds, as is the case in interference scenarios, this will be reflected in negative LD [12, 15, 26, 33]. To our knowledge, no analogue to this LD behavior has been found for adaptation in multiple (> 2) sites.

Here, we aim to extend these insights on the relationship between interference and LD to a context with multiple, but not infinite sites. First, we explore ways to generalize LD to multiple loci. *Multi-locus linkage dise-quibrium* (MLD) [34–37], has the advantage that it accounts for deviations from random association at more than two loci. MLD may thus appropriately reflect and characterize finite-sites interference. To compute MLD, we develop a recursive programming method applicable to up to seven loci. Second, we investigate the behavior of MLD under a finite-sites model of *multiple mutations interference* (MMI) [16]-a simplified model of interference-with parameter values calibrated to match HIV early infection dynamics. We focus on MMI due to its well-established theoretical framework [16] and its ability to appropriately describe more complex forms of interference [4].

We show that the evolution of MLD over time is interpretable and largely robust to drift. In deterministic scenarios, MMI causes MLD to oscillate, with a frequency proportional to the speed of evolution. Drift causes these features to become less pronounced, but still detectable. MLD oscillations can be further eroded by variation in fitness effects. We conclude that the wavelet-like oscillatory behavior of MLD results from, and is a robust signature of, finite-sites MMI.

## RESULTS

### Partition based definition makes MLD computationally tractable

To analyze how MLD is affected by multilocus interference during evolutionary dynamics, we first require a method to compute MLD. MLD, as formulated by Geiringer and Bennet, generalizes the notion of linkage disequilibrium from two to multiple loci using the principle that, due to the decay of allelic associations in haplo-types as a result of recombination, MLD between neutral genes should decrease exponentially over time [34, 35].

Consider *L* loci with alleles *i*_1_, *i*_2_,…,*i*_*L*_ and allele frequencies 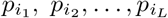
. Let 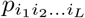, denote the frequency of haplotype **i** = *i*_1_*i*_2_ … *i*_*L*_, in the population. As introduced by Bennett [35], functions of allele and haplotype (i.e. gamete) frequencies, which satisfy the aforementioned decay condition are

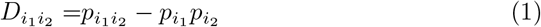

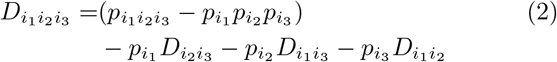

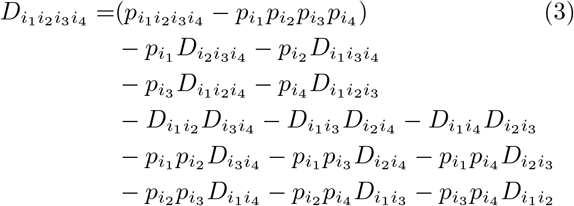

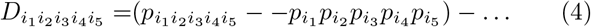

In equations (1–4), the terms 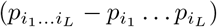 are called *Dausset’s disequilibrium* [38]. MLD, defined by 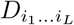, measures how much of Dausset’s disequilibrium cannot be attributed to lower-order associations of alleles. What remains is the unexplained over-or underrepresentation of the *L*^th^ order haplotype *i*_1_ … *i*_*L*_ only, or the *L*^th^ order MLD [34, 35]. Equations (1–4) are valid for multiple alleles at any locus *j*, but we will restrict our analysis to bi-allelic loci, *i*_*j*_ ∈ {0,1}.

Equations (1–4) for MLD can be expressed in a more concise fashion by means of partition theory, as shown by Gorelick and Laubichler [39, 40]. We add a superscript *L* to indicate the LD of *L*^th^ order, given *L* loci, and write:

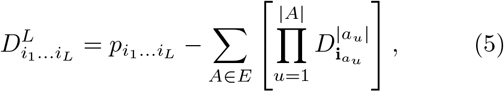

where E is the set partition of the set {1, …, *j*, …, *L*}, except for the trivial cell {{1, …, *L*}}. The *set partition* Ξ of a set *S* is a family of sets *A*, called *cells*, which contains all non-empty disjoint subsets of *S*, whose union is *S*. For example, the set partition Ξ of {1,2,3} is {{1, 2, 3}}, {{1, 2}, {3}}, {{1, 3}, {2}}, {{2, 3}, {1}} and {{1}, {2}, {3}}. The cell *A* = {{1, 3}, {2}}, has size |*A*| = 2, and elements *a*_1_ = {1, 3} and *a*_2_ = {2}.
 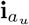 denotes a sub-haplotype: given for instance *a*_u_ = {1, 3}, then 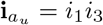. The disequilibrium of a single locus, 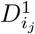, is defined as the allelic frequency 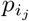 at that locus *j* [39].

Definition (5) allows disequilibria of higher order to be recursively defined in terms of disequilibria of lower orders. Recursive programming enabled us to computerize the algebra for higher order linkage disequilibria [41, 42] (see *Supporting Information, section 1, (SI.1)).* We obtained algebraic expressions for MLD, which depend only on haplotype and allele frequencies, for up to seven loci.

An alternative approach to MLD, due to Slatkin [43], defines it as the covariance between multiple alleles at multiple sites. The conclusions presented here apply to both definitions of MLD, although our analyses focus on the Geiringer-Bennet approach (see *SI.2*).

### Oscillations under the deterministic finite-sites approximation

We begin by describing the behavior of MLD under a simplified model of finite-sites interference, the *deterministic finite-sites MMI* (DFMMI) model, a deterministic analogue to MMI [16] in a finite-sites context. We retain MMI’s core assumption that all mutations will confer the same selective advantage, *s*, but remove stochasticity. Intuitively, selection moves the distribution of fitnesses of haplotypes steadily forward in the manner of a travelling wave [16, 44–46]; rapid growth of rare, fitter-than-average haplotypes expands the front of the distribution, while gradual loss of less-fit haplotypes contracts the distribution's tail [8, 10, 16]. Due to our interest in rapid adaptation of pathogens, simulation parameters were chosen to correspond to estimates from early HIV infection (see *SI.3*).

Our DFMMI simulations begin with a wildtype ancestor having a limited number *L* of possible beneficial mutations, which accumulate at a fixed rate; a rise in frequency of haplotypes with *k* mutations (*k*-mutants) is followed by a rise in frequency of *k* + 1-mutants every time period *τ*_inter_ (see *SI.3* for derivation), a constant independent of *k*. Within each *k*-mutant wave, we assume that the relative haplotype frequencies are equal. This assumption eliminates haplotype frequency imbalances stemming from genetic drift, allowing us to examine the dynamics of MLD in the absence of such complications. Moreover, it ensures that all possible 2^*L*^ haplotypes exist at some point in the evolutionary trajectory of the simulation - a *full escape graph* [47].

In each simulation, we allow a population to evolve for roughly 300 generations, calculating the *L*^th^ order MLD relative to the ancestral haplotype at fixed time intervals (see Figure 1). Unless otherwise noted, we subsequently refer to MLD relative to the ancestral haplotype, i.e. 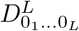, where *j* = 0_*j*_ denotes no mutation in the *j*^th^ allele.

In these DFMMI dynamics, the highest (*L*^th^) order MLD is initially zero. As single-mutant haplotypes appear and spread, the ancestral haplotype is outcompeted (Figs. 1A and B) and becomes under-represented relative to the expectation from random allelic associations. The MLD decreases during this process (Fig. 1C). During the remainder of the dynamics, the dominant *k*-mutants are replaced by successive *k* + 1 mutants (see Figs. 1B and 2A). The highest order MLD correspondingly oscillates from negative to positive. We found that the number of oscillations in the highest order MLD, *n*_*O*_, increases with the number of loci *L* simulated (*Figs. 3A* and C), andfollows the simple relationship:

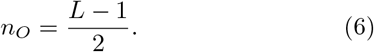

**FIG. 1.**
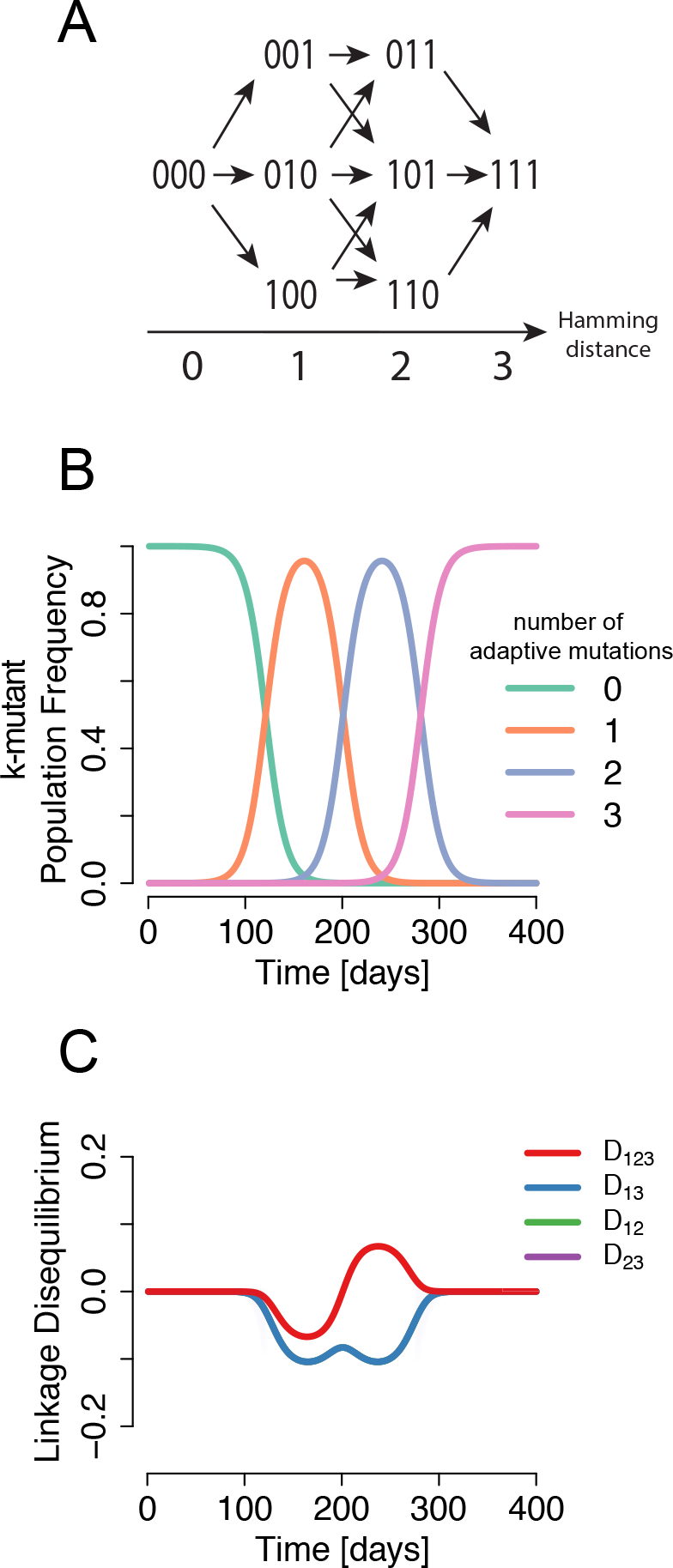
Origin of oscillations in multilocus linkage disequilibrium (MLD) A) The space of all possible haplotypes, starting from the wildtype (no mutations: all zeros). B) As evolution pushes the fitness distribution to higher Hamming distances, it generates a signature of over-representation of haplotypes with equal Hamming distance. This is reflected by the sequential rise and fall of k-mutant waves. C) Pairwise and three-locus Geiringer-Bennett linkage disequilibria, measured with the wildtype 000 as reference, over the course of the simulation (all the pairwise disequilibria overlap). When taken as a reference haplotype, all haplotypes with the same number of adaptive mutations produce an MLD of equal sign.

### MLD oscillations reflect DFMMI dynamics

The observed oscillations in the highest order MLD can be explained by the temporary dominance of *k*-mutant haplotypes in the population. In fact, the oscillations in highest order MLD reflect the acquisition of beneficial mutations. In the following, we refer to the highest order MLD simply as MLD.

As shown in Figure 1B, at any point during the dynamics, the population will consist mainly of haplotypes containing *k* mutants; i.e. *k*-mutant haplotypes will be over-represented. Therefore, the MLD relative to all *k*- mutant haplotypes will be positive. As mutation and selection push the population to higher fitness levels, *k* + 1-mutants spread. Then, the MLD relative to *k* + 1-mutant haplotypes will increase until it becomes positive.

A useful property of MLD in bi-allelic systems allows us to relate the MLD relative to a *k* mutant haplotype to the MLD relative to the ancestral haplotype: ∀*j*: *j* ∈ {1,…,*L*};

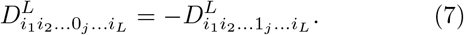

Equation (7) (see *SI*.2 for proof) can be interpreted as follows: if a reference haplotype is over-represented relative to our expectation, each haplotype with the opposite allele to the reference at a given locus must be equally under-represented.

Therefore, at any point during the dynamics, MLD relative to haplotypes containing a single beneficial allele (that is, single mutants) will be of equal magnitude, but opposite sign to MLD relative to the ancestral haplotype. Further, MLD relative to double-mutant haplotypes will be of equal magnitude, but opposite sign to MLD relative to single-mutant haplotypes; this also implies that MLD relative to double mutant haplotypes is equal to MLD relative to the ancestral haplotype. We conclude that when single or odd-*k* mutant haplotypes are over-represented (i.e. positive MLD), the MLD relative to the ancestral haplotype will be negative. In the same way, when double or even-*k* mutant haplotypes are over-represented, the MLD relative to the ancestral haplotype will be positive (see also *SI*.4, and Fig. S1).

Therefore, as the ‘traveling wave’ accrues subsequent beneficial mutations and the set of haplotypes that are over-represented (i.e., those haplotypes with positive MLD) shifts, the sign of the MLD relative to the ancestral haplotype also shifts. This explains both the observed oscillation in MLD and the relationship between the number of possible beneficial alleles and the number of observed oscillations (Eq (6)); there are *L* – 1 soft sweeps as additional beneficial mutations appear, and each sweep is reflected in a MLD half-oscillation.

The DFMMI model generates oscillations in MLD in another, rapidly evolving regime (when *τ*_inter_ is very small, see *SI*.5)). However, this particular MLD pattern is expected to be rare, and can be neglected when applying appropriate checks in data.

### Speed of evolution and MLD dynamics

As half-oscillations in MLD reflect partial sweeps of sequential layers of *k*-mutant haplotypes, we expect the frequencies of the MLD wavelets to correlate with the rate of evolution of the system. Let us assume that beneficial mutations accumulate at a stable rate, that is, that the population’s fitness wave proceeds at a well-defined constant speed *v* through fitness space. Then, the time for the fitness wave to accumulate one beneficial mutation corresponds to the time it takes for half an oscillation of the highest order MLD, *T/2,* where T is the MLD oscillation period. Thus, the speed of evolution of the fitness wave *v* must be related to the oscillation frequency of the MLD as follows:

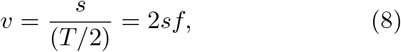

where *f* is the frequency of the oscillations.

### Retention of MLD oscillation properties under drift

Next, we tested whether MLD oscillations appear and can be detected and analyzed in the presence of drift, using a Wright-Fisher model with selection. To this end, we adapted the MMI model [16]: like MMI, our WF-model only considers drift-prone, beneficial mutations, each with the same effect, but our model considers beneficial mutations at finitely many loci. This model, previously employed in other studies [48, 49], is termed *finite-sites MMI* (FSMMI) model (see *SI*.6, and Fig. S2 for an example simulation).

As in the DFMMI model, our FSMMI framework and parameters are chosen to capture some features of early HIV within-host evolution, when HIV undergoes very rapid adaptation to the host environment [48, 49]. Specifically, we focus on regimes in which the population size is around *N* = 10^5^, the beneficial mutation rate per locus per generation is μ_b_ = 10^−4^ [33, 48–51], and each beneficial mutation carries the same selective advantage s between 0.01 − 0.3 [22, 51, 52]. The simulations are run with a population size *N* and selection acts on all loci from the start.

Unlike the DFMMI model described above, in FSMMI simulations beneficial mutations establish stochastically, breaking the symmetry in *k*-mutant haplotype frequencies. Thus, a full escape graph is not guaranteed. Despite this added stochasticity, beneficial mutations are still typically accrued in a sequential fashion (see Fig. 2), with subsequent *k*-mutants rising and falling in frequency. This is a prerequisite for MLD oscillations.

**FIG. 2.**
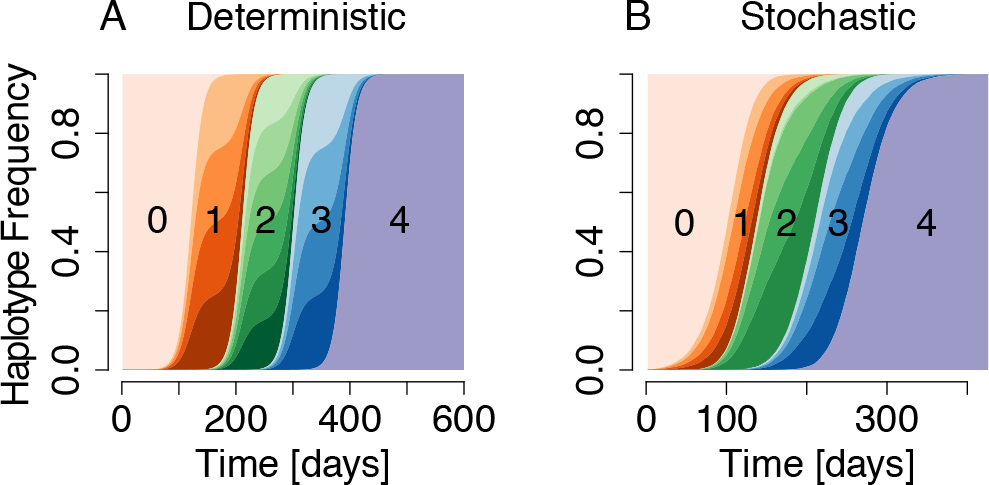
Haplotype dynamics of DFMMI and FSMMI simulations. A) Haplotype frequencies over the course of a DFMMI simulation with *L* = 4 loci. Beneficial mutations arise every *τ*_inter_ = 100 days (see *SI*.3) and begin to sweep at a rate *ϵ* = 0.095 (see *SI*.3, eqn. (S11)). Colors indicate haplotypes with an equal number of mutations *k*. B) Haplo-type frequencies over the course of a simulation of the FSMMI model with *L* = 4 selected loci, selection coefficients per mutation *s* = 0.1, population size *N* = 10^5^ and beneficial mutation rate *μ*_*b*_ = 10^−4^ per locus per generation.

In fact, both wavelet-based statistical tests and Fisher tests for hidden periodicities indicate that oscillations in the highest order MLD persist under FSMMI (see *SI*.7, Figs. S3 and S4). However, as expected, the oscillations tend to be less precise than in the DFMMI case (Fig. 3A vs 3D) and *n_*O*_* full oscillations are not always realized. This dampening of signal is likely due to portions of the haplotype space remaining unexplored in stochastic simulations.

**FIG. 3.**
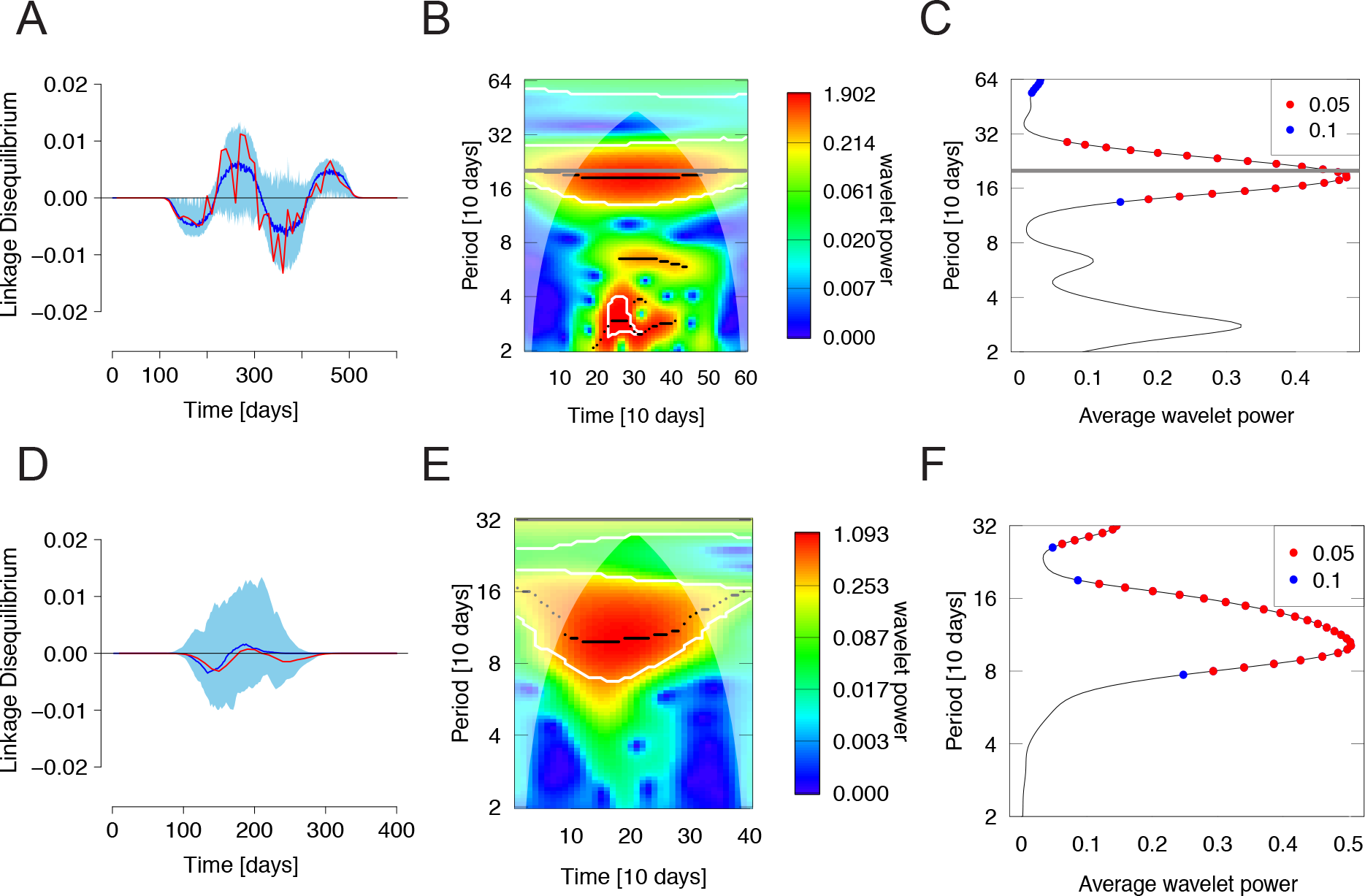
MMI-induced MLD oscillations are still detectable under drift. A) Oscillation of the fifth order MLD in a symmetric full escape graph. The dark blue line is the median of a set of 200 runs, and the upper and lower bounds of the light blue area represent the 2.5 and 97.5 percentiles of all measured LDs. The MLD was calculated every 10 days using a sample size of 20 haplotypes. The red trajectory represents the measured LD from one particular repeat. B) The wavelet power spectrum in the time-period domain of the fifth order MLD values obtained with the sampling points of the red line in A) [53, 54]. The horizontal grey line is the true oscillation period of the red time series in A). The white contour lines indicate regions where the power spectrum values are significantly (< 5%) non-random. The black lines indicate local power spectrum maxima. The half-transparent region demarcates a low-confidence wavelet power region. C) The time-averaged wavelet power spectrum. The red and blue dots indicate whether the null-hypothesis that the the time-averaged wavelet power may have been generated by white noise is rejected at below 0.05 or 0.1 significance levels, respectively. The maximum spectral density is attained close to the simulated period of *T* = 200 days (horizontal thick grey line) of the oscillations. D) The analogous situation to A) for 100 simulation runs of the FSMMI model with selection, run with parameters *L* = 4, *N* = 105, *μ*_*b*_ = 10*−4* and *s* = 0.1. Samples are taken every 5 generations or 10 days. E) Wavelet power spectrum of one randomly chosen MLD trajectory (red line in D)). F) Analogous to C), but without the horizontal line indicating expected value.

We further investigated whether the frequencies of these MLD signals may be estimated. To this end, we computed the wavelet power spectrum [53-55], of the simulated dynamics (Fig. 3B,3E) and used it to infer the frequency at which MLD oscillates (see *Materials and Methods*). As shown in Fig. 3C for a DFMMI benchmark, even in stochastic simulations (Fig. 3F), wavelet analysis can confidently reconstruct the frequency of MLD oscillations.

We proceeded to examine whether the MLD oscillations under FSMMI also retain other features displayed under DFMMI, such as equation (8). To this end, we compared the rate of evolution estimated using the MLD oscillation frequency, 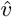, to the rate of evolution expected under infinite-sites MMI theory, *v*_MMI_ [16] (see Fig. 4, *SI*.8). *v*_MMI_ serves as benchmark because there does not exist a clear ground truth for the speed of evolution under finite-sites.

**FIG. 4.**
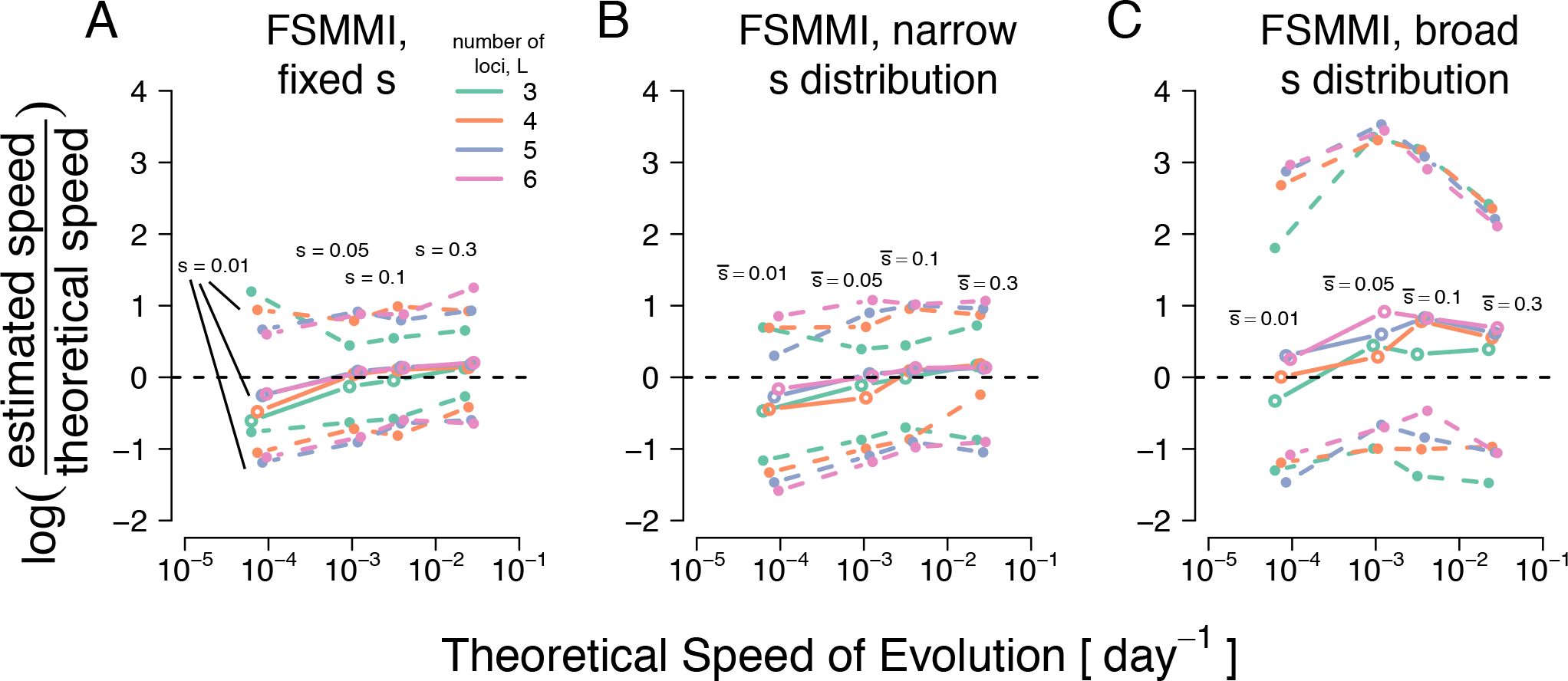
Estimates of speed of evolution based on MLD-oscillations versus MMI theory [16]. A) Estimates of the speed of evolution 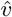 obtained by wavelet analysis for FSMMI model simulations run for *L* = 3, 4, 5 and 6 loci, and selection coeficients *s* ∈ {0:01, 0:05, 0:1, 0:3} with population size *N* = 10^5^. *v* is estimated with equation (8) (*s* is known), where for *f* we use the MLD-based oscillation frequency estimate (Fig.3). The inter-sampling period was Δ*t* = 2 days. Colored open circles and filled circles correspond to medians and nonparametric confidence intervals (95%), respectively, from those simulations among 100 runs that displayed significant (< 0:05 level for wavelet-based test) oscillations. B) Analogous figure for narrow FSMMI model. Here, to compute the speed of evolution *v* from the inferred oscillation frequency *f* we use the average 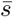 of the Gamma distribution from which the selection coeficients were sampled (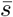 values equal to s values in FSMMI). C) Analgous to B), but for the broad FSMMI model.

Figure 4A shows that the rate of evolution inferred by MLD dynamics from our simulated FSMMI model, 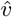 from (8) with *N* = 10^5^ is close to *v*_MMI_, the predicted rate of evolution in infinite-sites MMI theory, [16].

When *N* varies from 10^3^ to 10^6^, the mismatch between 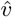 and *v*_MMI_ first decreases, and then begins to increase again (see *SI* Fig. S5, left column). As expected, when population size values fall to 10^3^ or lower, the interference effects fade [16], and the 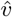-to-*v*_MMI_ differences increase markedly. For populations sizes around 10^6^, 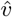 becomes smaller than *v*_MMI_, but remains within confidence intervals for almost all studied cases.

We performed an analogous test using Crow-Kimura-Felsentein (CKF) theory, *v*_CKF_ [56] (see *SI* Fig. S6, left column). Apart from a systematic positive bias of 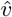 relative to *v*_CKF_, the above patterns are largely retained.

### Retention of MLD oscillation properties under drift and dissimilar fitness effects

Despite its usefulness for mathematical analysis, the assumption that all loci have identical fitness effects is unlikely to be perfectly satisfied in biological systems. To address this issue, we devised two further models, *narrow FSMMI* and *broad FSMMI* (see *Materials and Methods*). In the narrow FSMMI model, the selective coefficients *s*_*j*_ of a mutated locus *j* are drawn from a Gamma distribution with average 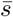. The broad FSMMI model adopts a distribution of fitness effects (DFE) which falls off in an over-exponential manner [57]. Again, the average *s*_*j*_ is predefined.

We observed that narrow FSMMI simulations produced detectable oscillations nearly as often as FSMMI (see *SI*7, Figs. S7). Significant signals of MLD oscillation were also often detected under broad FSMMI (see *SI*7, Figs. S8). However, due to exacerbated symmetry breaking of the escape graph, these were substantially less frequent than in the simulations under narrow FSMMI. Further analysis is concentrated on simulations with non-random MLD time series behavior.

Figure 4B shows that the narrow FSMMI model leads to MLD-based estimates similar to the those from the FSMMI model. whereas broad-FSMMI estimates (4C) deviate more strongly from theory. Confidence intervals around estimates broaden as selective coefficients tend to become more dissimilar. Varying *N* affects narrow and broad FSMMI estimates similarly as it does FSMMI estimates (see *SI* Fig. S5). A similar picture emerges when using CKF-theory as benchmark (see *SI* Fig. S6).

To further corroborate these results, we also compared 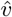 values with estimates from alternative methods of inference of the speed of evolution (see *SI*9). The performance of MLD-based estimates relative to other, reasonable estimates of the speed of evolution 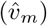 is generally very similar to their performance relative to predictions from MMI theory (see *SI,* Fig. S9, and S10 for a systematically biased estimator).

Lastly, as the assumptions of infinite-site MMI theory are not met in our FSMMI simulations, we assessed *v*_MMI_’s appropriateness as a benchmark. To do this, we compared *v*_MMI_ against *v*_m_. The match between MMI theory and empirical estimates is generally good (see *SI*, Fig. S11), suggesting that the infinite-sites assumption in MMI theory does not adversely affect its predictive utility for finite-sites systems within the parameter ranges here studied. This justifies the use of MMI theory as a benchmark for wavelet-based estimates.

## DISCUSSION

We have developed new computational tools to calculate multilocus linkage disequilibrium (MLD), a statistic that quantifies the nonrandomness of allelic associations across loci, accounting for contributions to haplotype structure stemming from subgroups of loci. We show that, in simulated deterministic haplotype dynamics with (i) rapid accrual of a finite number of strongly beneficial mutations with similar fitness effects and (ii) tight linkage between loci (i.e. a MMI regime), MLD dynamics display a wavelet-like temporal pattern. We find that these oscillations can be explained by successive sweeps by haplotypes containing increasing numbers of beneficial mutations in combination with specific mathematical properties of MLD expressed in Eq. 7. We demonstrate that the frequency of these oscillations is proportional to the rate of evolution. Finally, we show these oscillations are robust to evolutionary stochasticity and some degree of variation in the fitness effects of mutations. However, these properties are gradually lost as the fitness effects become more dissimilar. Thus, MLD dynamics may contain information relevant to the study of the short-term evolution of microorganisms under very strong selection, including human pathogens such as HIV, in which a finite number of loci experience strong selection.

Moreover, the detection of MLD oscillations depends on accurate haplotype frequency estimates, not obtained in most within-host evolution studies, and in HIV in particular. However, the continuous improvement of sequencing technologies is likely to allow for deep and dense sampling in the future, producing appropriate datasets.

While the MLD behavior we describe exhibits important evolutionary phenomena, the wavelet-based approach we present for inferring the speed of evolution will likely be inefficient in natural populations. Rather than providing a new estimation method, this study aims to elucidate how evolutionary dynamics are simultaneously manifest in both haplotype frequency dynamics and multilocus linkage disequilibria. This is a necessary first step before the MLD perspective on evolutionary dynamics can offer broader applicability.

Further, our method currently ignores the role of epistasis. In escape mutations of HIV, the pathogen which inspired this work, we are unaware of evidence for epistatic interactions. However, other intragenic mutations are likely to give rise to epistasis [58–61]. If epistasis dominates over selection (sign epistasis), the evolutionary dynamics are likely to halt at a local or global fitness peak (i.e. not the full escape haplotype) [62]. Then, at mutation-selection balance, an MLD signal should be maintained that is constant and not oscillatory. Extensions of this work may thus help to differentiate epistasis-dominated from weak-or no-epistasis scenarios.

We also assume that loci under selection are readily detectable. This is true in the case of epitopes targeted by cytotoxic T lymphocytes in HIV-infected patients (see [25]), but may not be true elsewhere. We did not investigate the scenario where only *L*′ are tracked, but where *L* > *L*′ are under selection. Unknown loci may exacerbate escape graphs’ asymmetries, thereby further dampening any MLD signal. Further work is needed to fully ascertain the impact of these effects.

Another benefit of our approach to interference is that it draws from an underexplored perspective on evolution that considers the role of linkage disequilibria, and its important statistical inference machinery. In fact, very little use has been made of MLD in the context of population genetics, in particular the study of interference [32]. This may be due to different definitions of linkage disequilibrium at multiple loci [34, 35, 43, 63, 64]. The crucial advantage of the Geiringer-Bennet MLD is that its maximum likelihood estimate always exists [65], a very useful property for estimation.

The other central benefit is MLD’s capacity to characterize a population as evolving under MMI. Most simply, the presence of MLD oscillations of the type described here suggests that the population under study is evolving under an MMI regime. MMI occurs in populations with specific characteristics; namely (i) a large supply of beneficial mutations [16] (ii) beneficial mutations that confer similar, strong selective advantages [16], and (iii) low enough recombination rates that beneficial mutations are likely to compete rather than recombine onto a single haplotype. Therefore, observed MLD oscillations provide valuable information with respect to these critical population genetic parameters.

## MATERIALS AND METHODS

### Oscillation estimation by means of signal processing techniques

To identify oscillations of MLD in the simulation data, we developed a detection scheme based on wavelet analysis. For each run, we calculated the highest order linkage disequilibrium at each of *M*_*s*_ sample points from the sampled data, that is, *M*_*s*_ MLD-values {*x*_*n*_}, where *n* ∈ {0,…, *M*_*s*_ − 1}. Sample data *x*_*n*_ are assumed to have been obtained at constant inter-sampling periods *dt*, and can be expressed as a vector x with entries *x*_*n*_.

We analyzed the wavelet power spectrum of x (Repackage *WaveletComp,* [66]). An oscillating LD measure of *L* loci will maximally generate *L* − 1 half-oscillations, starting with a negative half-oscillation. Even if damped, such wavelet-like oscillations should leave traces in the frequency spectrum that are close to the frequency of a full period, *T*.

The *wavelet transform* of the data x is given by:

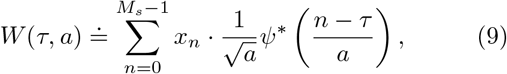

where

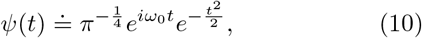

is the Morlet wavelet. *τ* is called the *translation time,* whereas *a* is termed *scaling factor.* The superscript * denotes the complex conjugate. The nondimensional frequency *ω*_0_ is set to 6, such that the scale *a* becomes almost identical to the Fourier period.

To compute all values of *W*(*τ*, *a*) and its derivates, we used the package *WaveletComp* [66]. The wavelet transform is computed over a standard set of values of *τ* and *a*. *τ* is varied from 0 to *M*_*s*_ − 1; that is, by multiples of the time increment *dt*. The scaling factors *a* determine the coverage in the period domain. They are set to vary as *a*_min_. 2^*j.dj*^, with *j* = 0, …, *J* and where *a*_min_ = 2 is the minimum scaling factor used, *dj* = 1/20 is the number of steps per analyzed octave and the maximum period analyzed, *a*_min_. 2^*J.dj*^, is 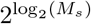(which determines *J*).

The *wavelet power spectrum* is defined by:

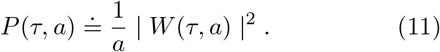

The value of P(*τ*, *a*) at a coordinate pair (*τ*, *a*) serves as a measure of confidence that the time series x is oscillating at a frequency corresponding to *a* at translation time *τ*.

To measure the frequency *f* of an MLD time series, we first identify the value pair (*τ*_max_, *a*_*w*,max_) for which *P*(*τ*, *a*) was maximal. A p-value for the null-hypothesis that there is no periodicity in x is provided by *Wavelet-Comp*, using a statistical test based on the work of Cazelles et al. [53–55].

To reduce computational burden, the time series {*x*_*n*_} was trimmed for analysis. Simulation runs were all performed over the simulation time of 2000 generations. However, since *L*-mutants frequently fix well before 2000 generations have elapsed, most late *x*_*n*_ values are zero. To speed up computations, we identified the last non-zero *x*_*n*_ value for each series, *x*_*z*_, and replaced the series {*x*_*n*_} by {*x*_0_,…, *x*_*z*_, 0,…, 0}, concatenating 20 zero values to the end of the series. This modified series was then used for further analysis.

When stochastic simulations were run, MLD could be zero for the entire time course at low population sizes and very different selection coefficients. In these cases, the oscillation frequencies were set to correspond to zero. For Fisher's hidden periodicities test the p-values for the null that no periodicities exist in the all-zero signal were set to unity.

### Distribution of fitness effects employed in finite-sites MMI models

In this study, three different distributions of fitness effects were used to run Wright-Fisher simulations with a finite number of loci to examine the robustness of (8). The first set of simulations were run with the simplest possible assumption: all loci confer the same selective advantages.

The second set of simulations were run with selection coefficients drawn from a Gamma distribution:

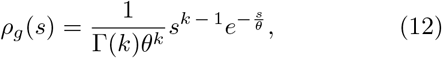

where Γ is the Gamma function, *k* is the shape parameter and *θ* is the scale parameter. For our simulations, we used the values *k* = 400, and *θ* = {7.5 . 10^−4^, 2.5 . 10^−4^, 1.25 . 10^−4^, 2.5 . 10^−5^}. The average s under that distribution is 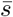 = *kθ*. The variance is always set to 10% of 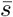.

The third set of simulations were run by drawing the selection coefficients from an exponential-like distribution of fitness effects with parameters that favour MMI conditions [49, 57]. Specifically, the distribution of fitness effects used was:

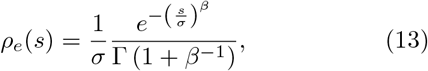

where *β* is a steepness parameter that indicates whether the distribution follows an over or under-exponential decline as *s* increases, and σ roughly corresponds to the inverse of the rate parameter in an exponential distribution. The average selection coefficient sampled, 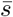, is given by σΓ(2/*β*)/Γ(1/*β*). When *β* is one, *ρ*_*e*_(*s*) is exponentially distributed. In this study, we used parameters for average values of s that are *β* = 1.4 and σ = {0.432,0.144,0.072,0.0144}.

## ACKNOWLEDGMENTS

The authors thank Fabio Zanini, Roland Regoes, Frederic Bertels, Massimo Maiolo and the Feldman lab members for stimulating discussions. This work was supported by by the Swiss National Science Foundation (grant number P2EZP3_162257 to VG). AH was supported in part by The Stanford Center for Computational, Evolutionary and Human Genomics (CEHG) doctoral fellowship at Stanford. MWF was supported in part by CEHG, and by the Morrison Institute for Population and Resource Studies.

